# Programmed cell death and the origin of wing polyphenism in ants: implications for major evolutionary transitions in individuality

**DOI:** 10.1101/2024.02.14.580404

**Authors:** Lisa Hanna, Brendon E. Boudinot, Jürgen Liebig, Ehab Abouheif

## Abstract

Major evolutionary transitions in individuality occur when solitary individuals unite to form a single replicating organism with a division of labor between constituent individuals. Key examples include the evolution of multicellularity, eusociality, and obligate endosymbiosis. Programmed Cell Death (PCD) has been proposed to play an important role during major transitions to multicellularity, yet it remains unclear to what extent PCD plays a role in other major transitions. Here we test if PCD was involved in the major transition to eusociality in ants, where solitary individuals united to form eusocial colonies with a division of labor between a winged queen caste and a wingless worker caste. The development of wings in queens but not in workers in response to environmental cues is called wing polyphenism, which evolved once and is a general feature of ants. Both wing polyphenism and eusociality evolved at the same time during the origin of ants and were likely intimately linked––the suppression of wings in workers may have reduced their ability to participate in mating flights thereby reinforcing the reproductive division of labor within the parental nest. We therefore tested whether PCD plays a role in the degeneration of wings during development of the worker caste across the ant phylogeny encompassing species with both ancestral-like and derived characteristics. We show that PCD, mediated by the apoptosis pathway, is present in the degenerating wing primordia of worker larvae in 15 out of the 16 species tested. Using ancestral state reconstruction, we infer a role for PCD in regulating wing polyphenism in the last common ancestor of all extant ants. Our findings provide evidence that a degenerative mechanism (PCD) plays a role in the origin of wing polyphenism, and therefore, in facilitating the major transition to eusociality in ants. PCD may generally play a key role in the evolution of biological complexity by facilitating major transitions at different scales, such as multicellularity and eusociality.

## INTRODUCTION

Major evolutionary transitions in individuality occur when solitary individuals unite to form a single replicating individual. During the history of life, such transitions include the integration of solitary prokaryotic cells to form the eukaryotic cell, the integration of single-celled organisms to form multicellular organisms, the integration of solitary organisms to form eusocial colonies, and the integration of distantly-related organisms to form obligate endosymbioses (Buss, 2014; Maynard-Smith & Szathmary, 1997; West et al., 2015). Despite great advances made in understanding the evolutionary principles underlying these major transitions, the developmental mechanisms that facilitated these events remain poorly explored.

Recent progress in understanding the major transitions to multicellularity have led to the proposal that programmed cell death (PCD) is a mechanism that may have played an essential and general role in the emergence of major evolutionary transitions (Durand et al., 2019; Durand et al., 2016; Huettenbrenner et al., 2003). Unlike other modes of cell death, PCD is a genetically regulated process that occurs under specific physiological conditions (Galluzzi et al., 2018). It has ancient origins, and in addition to multicellular life, is frequently found in unicellular eukaryotes and prokaryotes (Ameisen, 2002). PCD has been proposed to play a role in facilitating major transitions through the formation of alternative phenotypes, and therefore, the formation of a reproductive and/or morphological division of labor (Durand et al., 2019; Huettenbrenner et al., 2003; Libby & Ratcliff, 2014). For example, in *Dictyostellium*, PCD facilitates the formation of two alternative cell types under stressful environments, in which one cell type undergoes programmed cell death becoming stalk cells, thereby helping the other cell type to disperse their spores (Arnoult et al., 2001). Another example includes the experimental evolution of multicellularity in a unicellular yeast, which led to the formation of alternative cell types within the newly evolved multicellular unit (called snowflakes). Some cells undergo PCD allowing the evolved multicellular snowflakes to fracture and form clusters that can propagate and disperse (Libby & Ratcliff, 2014; Ratcliff et al., 2012). Finally, in the eusocial honeybee, PCD facilitated the formation of a reproductive division of labour between the queen and worker castes, such that PCD reduces ovary development in worker-destined larvae but not in queen-destined larvae (Durand et al., 2019; Hartfelder & Steinbruck, 1997). Although these examples are consistent with the hypothesis that PCD may facilitate the origin of major evolutionary transitions, to our knowledge, it has not been formally tested using phylogenetic comparative methods with a sampling of ancestral and derived species within a taxonomic group.

Here we test the hypothesis that PCD contributed to the major evolutionary transition to eusociality in ants (family Formicidae) (Boudinot, Richter, et al., 2022; Holldobler & Wilson, 1990). Approximately 150 million years ago, solitary wasp-like individuals united to form the first eusocial ant colonies. Fossil and phylogenetic evidence shows that these ancestral ant colonies possessed a wing polyphenism between the queen and worker castes (Boudinot, Khouri, et al., 2022; Hanna & Abouheif, 2021). Wing polyphenism is the ability of an egg to develop either into a winged queen or wingless worker in response to environmental cues. Current data suggests that nutrition, social interactions, and temperature are key environmental cues, which are mediated by Juvenile Hormone (JH) to regulate the developmental switch between winged queens and wingless workers (Hanna & Abouheif, 2021; Penick et al., 2012; Wheeler, 1986) (Fig. 1a). Typically, adult queens and males use their wings to participate in nuptial flights, while the wingless worker caste engages in foraging and brood care on or under the ground (Holldobler & Wilson, 1990). The origin of eusociality and wing polyphenism in ants appeared during the same time period and are now universal traits (excepting for a few secondary losses) (Boudinot, Khouri, et al., 2022; Boudinot, Richter, et al., 2022; Hanna & Abouheif, 2021; Peeters, 2012; Perrichot et al., 2008). It has therefore been proposed that eusociality and wing polyphenism are intimately linked: the origin of eusociality may have facilitated the origin of wing polyphenism, or alternatively, the origin of wing polyphenism may have facilitated the origin of eusociality (Hanna & Abouheif, 2021). In the early lineages of ants, queens and workers were inferred to have similar reproductive capacities, such that they are able to mate and store sperm, and therefore, reproductive division of labor primarily occurs through behavioral regulation in these lineages (Peeters, 1991). Consequently, the origin of wing polyphenism may have played an important role in the origin of eusociality by limiting worker’s participation in nuptial flights and dispersal from the parental nest. This in turn limited the ability of workers to mate and reproduce thereby reinforcing the reproductive division of labor in the early lineages of ants (Bourke, 2013; Hanna & Abouheif, 2021; Khila & Abouheif, 2008; Nowak et al., 2010). In support of this inference–– both ants and termites are the only two groups of insects where all species are eusocial, and all species have a wing polyphenism (Wilson, 1971). We therefore ask if PCD contributed to a major evolutionary transition in eusociality in ants by playing a key role in regulating wing polyphenism by degenerating wings in the worker caste during development.

**Figure 1.**
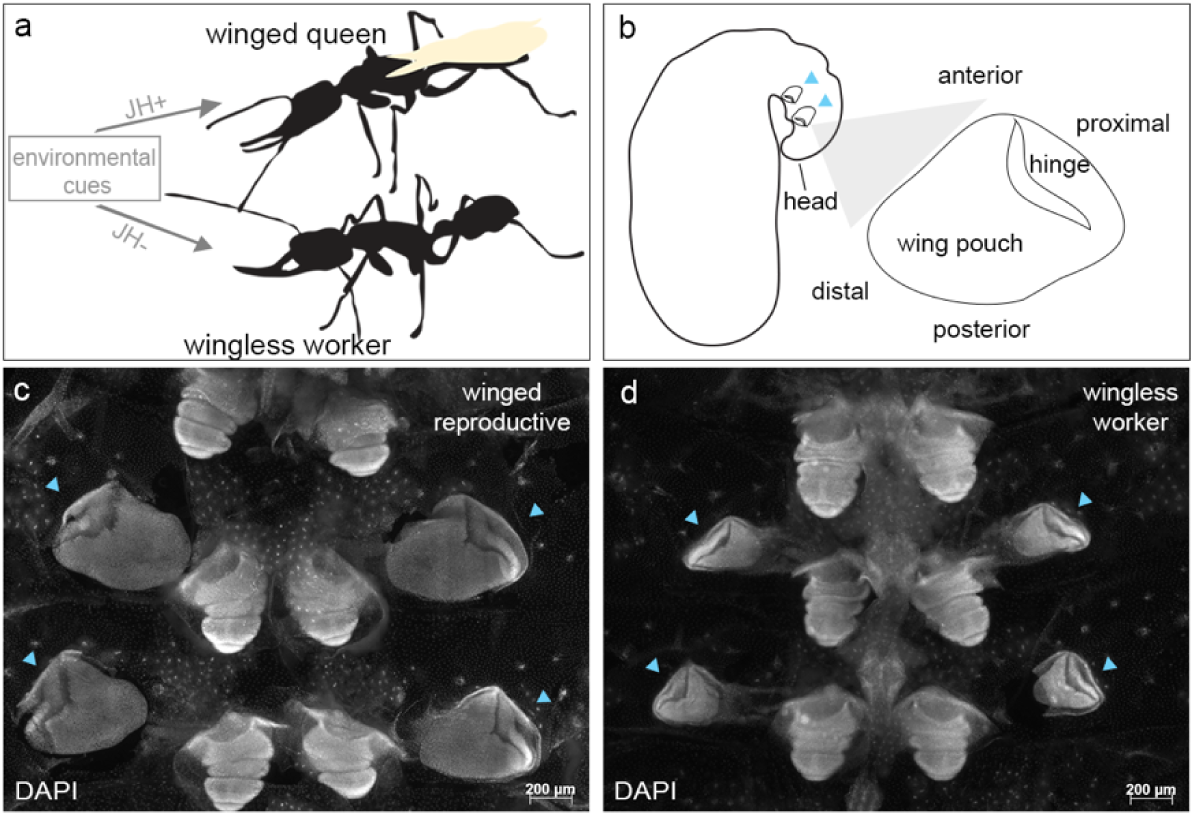
Wing polyphenism in the ponerine ant *H.saltator*: (a) environmental cues determine the development of winged queens (wings indicated in yellow) or wingless workers and reproductive workers ‘gamergates’ (not shown). (b) a cartoon illustration of *H. saltator* larva during terminal development indicating the location of the wing discs (blue arrowheads) and the general axes and regions of the wing disc in ants. Dissected terminal stage larvae in *H. saltator* stained with DAPI showing: (c) wing discs in winged males and (d) rudimentary wing discs in wingless female workers (blue arrowheads).

Individuals in the worker caste of ants, although completely wingless as adults, transiently develop wing discs during larval development similar to the winged caste (Fig. 1b–d, blue arrowheads) (Abouheif & Wray, 2002; Dewitz, 1878; Rajakumar et al., 2018; Wheeler & Nijhout, 1981). These wing discs are therefore considered as rudimentary organs, which Hall (2003) defines as an embryonic primordium of a more fully developed feature (the insect wing) found in an ancestor (a winged solitary wasp). In several derived ant species, the winged castes (queens and males) deploy the highly conserved wing gene regulatory network (GRN) to specify and regulate wing development. In contrast, the wing GRN is interrupted to halt development of wing rudiments in the wingless worker caste (Abouheif & Wray, 2002; Shbailat & Abouheif, 2013; Shbailat et al., 2010). For example, in the derived ant genus *Pheidole*, the worker caste is comprised of two types of workers––the small-headed minor workers and big-headed soldiers. Only soldier-destined larvae initiate the development of a single pair of large rudimentary forewing discs. Although GRN expression is conserved in the winged castes, it is interrupted in these soldier rudimentary forewing discs (Abouheif & Wray, 2002). Subsequent studies, however, showed that apoptosis (the most common genetic pathway of PCD) also plays a key role in eliminating these rudimentary forewing discs (Sameshima et al., 2004; Shbailat et al., 2010). Specifically, apoptosis first appears in the hinge of these discs (Fig. 1b) during the last stages of larval development and then begins to spread to the wing pouch, the peripodial membrane (an integral part of wing imaginal disc), and the tissue that attaches the disc to the cuticular wall (Sameshima et al., 2004; Shbailat et al., 2010). At this point, the soldier rudimentary forewing discs show clear signs of degradation and decrease in size. In contrast, no apoptosis is detectable in the wing discs of winged castes (males and queens) during all stages of larval development. This means that both GRN interruption points and apoptosis play a role in halting wing development in the wingless worker caste in *Pheidole.* In contrast to *Pheidole* and other derived ant species, no wing GRN interruptions were detected in the wingless workers in two ponerine species from the genus *Mystrium*, which are thought to reflect the ancestral characteristics of ants (Behague et al., 2018). This lead Behague et al (2018) to propose that apoptosis may have been the principal mechanism co-opted to eliminate wing discs in the wingless worker caste in this lineage of ants. However, at present, the only evidence that apoptosis plays a role in regulating wing polyphenism in ants comes from a single ant genus *Pheidole*. It therefore remains unclear how phylogenetically widespread this role of apoptosis is and whether it played a role in the origin of wing polyphenism.

Therefore, the main goal of our study is to test whether apoptosis played a role during the major transition to eusociality by testing its role in the origin of wing polyphenism in ants. To test this hypothesis, we used phylogenetic comparative analyses focusing on the two major clades of ants: the ‘poneroid’ and ‘formicoid’ clades. Species in the poneroid clade, like in the genus *Mystrium* we discussed above, are of key importance to understanding the mechanism that may have been present in the early history of wing polyphenism, since these species exhibit several characteristics that are generally considered ancestral within ants (Burchill & Moreau, 2016; Keller & Peeters, 2022; Peeters, 1997; Wilson & Holldobler, 2005). Some of these characteristics include low queen-worker dimorphism, small colony size, predatory behavior, and similar reproductive capacity between queens and workers (Keller & Peeters, 2022; Wilson & Holldobler, 2005). In contrast, species in the formicoid clade, like in *Pheidole*, largely exhibit derived social characteristics, such as high queen-worker dimorphism, large colony size, the evolution of worker subcastes, and large differences in reproductive capacity between queens and workers (Holldobler & Wilson, 1990; Khila & Abouheif, 2008; Matte & LeBoeuf, 2024; Peeters, 1991; Wills et al., 2018; Wilson, 1953). Therefore, we first tested for the presence of apoptosis in the winged males and wingless workers and across several developmental stages of a species belonging to the poneroid clade *Harpegnathos saltator* of, which is thought to have ancestral (plesiomorphic) characteristics. We then tested for the presence of apoptosis in an additional 15 species across the ant phylogeny, belonging to both the poneroid and formicoid clades. Finally, we use our developmental data to conduct an ancestral state reconstruction analysis to test whether apoptosis was present during the origin of wing polyphenism.

## RESULTS

In *H. saltator*, a species with ancestral-like characteristics, we first tested for apoptosis in the wing discs of male larvae that develop into adults with fully developed wings. We found that during the early and mid-periods of the last larval instar, the male wing discs show some small clusters of apoptosis signal (Fig. 2a–d and 2a’–d’, white arrowheads). However, by the larval-pupal transition, the apoptosis signal is no longer present (Fig. 2e, e’). In contrast, in the rudimentary wing discs of worker larvae, apoptosis first appears in the wing hinge and pouch during the early period of the last larval instar (Fig. 2f, f’ and 2g, g’, white arrowheads). By the mid-period, the apoptosis signal expands within the hinge, wing pouch, the peripodial membrane, and the tissue attaching the disc to the cuticle (Fig. 2h, h’ and 2i, i’). Finally, by the larval-pupal transition, apoptosis spreads throughout the entire rudimentary wing disc, which at this stage is highly reduced in size (Fig. 2j, j’). Together, our results show that apoptosis plays a role in eliminating the rudimentary wing discs in wingless workers, but not the wing discs in the winged castes in *H. saltator*.

**Figure 2.**
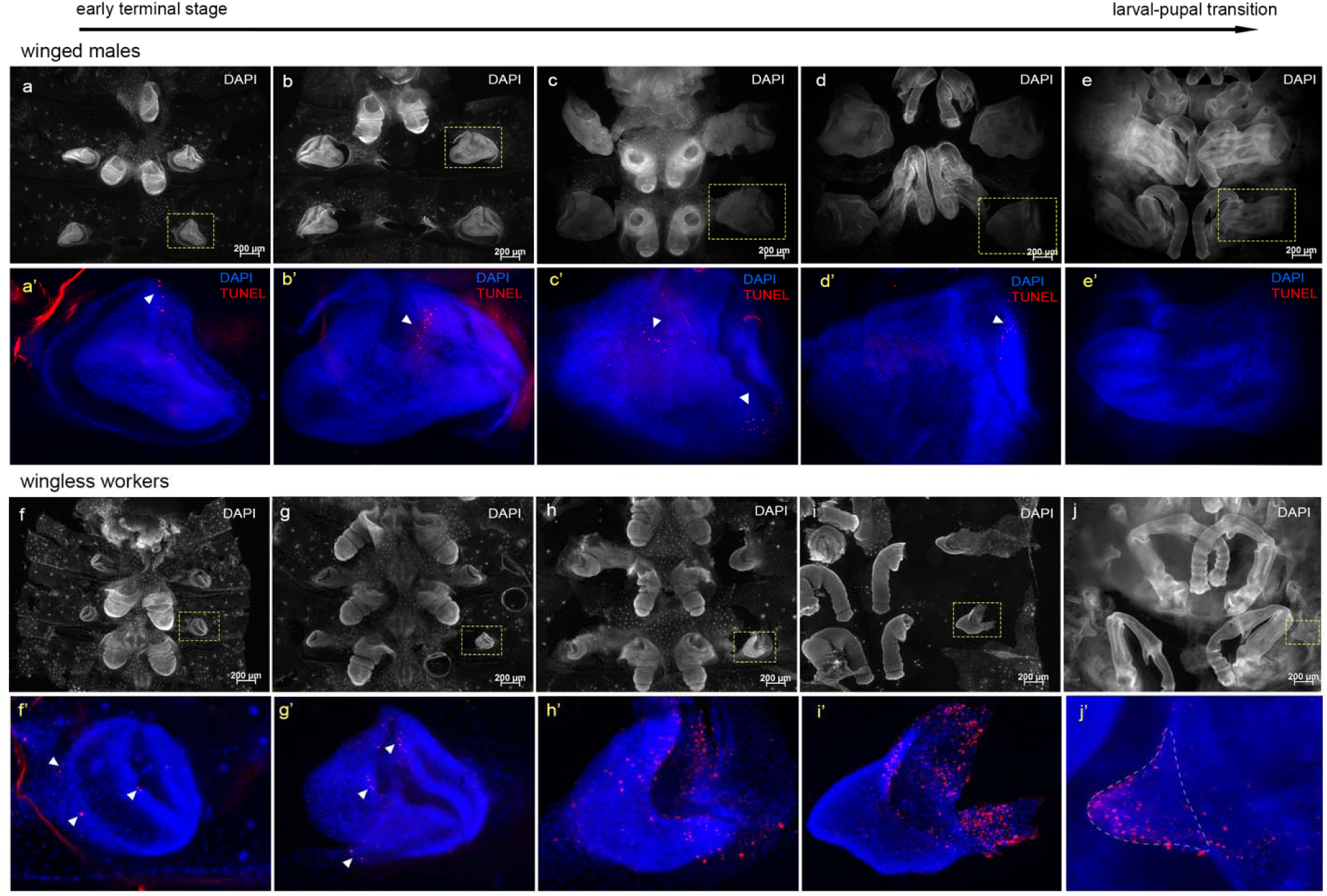
Programmed cell death in the wing discs of wingless workers and winged male larvae of *H.saltator*. (a–e) Larval developmental series of winged males from early/mid terminal stage to larval-pupal transition stained with DAPI. (a’–e’) a magnified view of a wing disc from the larva in the panel directly above showing TUNEL signal (red) and DAPI (blue). White arrowheads in (a’–d’) indicate areas with TUNEL expression. (f–j) DAPI staining of a larval developmental series of wingless workers during terminal larval development corresponding in stage to those shown in the males. (f’–j’) a magnified view of a wing disc from the larva in the panel directly above showing TUNEL signal (red) and DAPI (blue). White arrowheads in (f’, g’) indicate regions where TUNEL is expressed. TUNEL expression in (h’–j’) is widespread. Dashed line in (j’) outlines the wing disc area. Yellow dashed boxes in (a–e) and (f–j) indicate the wing disc magnified in the panels below. Images in panels (a–e) and (f–j) are to scale. Images in panels (a’–e’) and (f’–j’) are not to scale. All wing discs are oriented with distal axis on the left and proximal axis on the right (see Fig. 1b).

We next tested for apoptosis in the rudimentary wing discs of worker-destined larvae across 15 additional species from five major subfamilies of ants (Fig. 3; Supp. 1). We first tested two species belonging to the poneroid clade: *Odontomachus brunneus* (Ponerinae) and *Stigmatomma pallipes* (Amblyoponinae). Similar to *H. saltator*, apoptosis in the rudimentary wing discs of these ponerines first appears during the early to mid-period of the last larval instar and then spreads throughout the rudimentary wing disc (Fig. 3a, a’ and 3b, b). We next tested 13 species from the formicoid clade. Despite the great variation in size and shape of rudimentary wing discs across species in this clade, we found that in the early-mid period of the last larval instar, apoptosis is either absent (Fig. 3c, f, and i), or generally initiates in the hinge region (Fig. 3d, e, g, h, j, and Supp.1). During the later period of the last larval instar, the signal spreads throughout the disc (Fig. 3d’, e’, g’, h’, i’, j’, and Supp. 1). *Tetramorium immigrans* (Myrmicinae), which possesses very small rudimentary wing discs, is the only species where we do not observe apoptosis signal during any stage of the last larval instar (Fig. 3f, and f’). Our results show that the presence of apoptosis in the rudimentary wing discs of wingless workers is widespread across ants.

**Figure 3.**
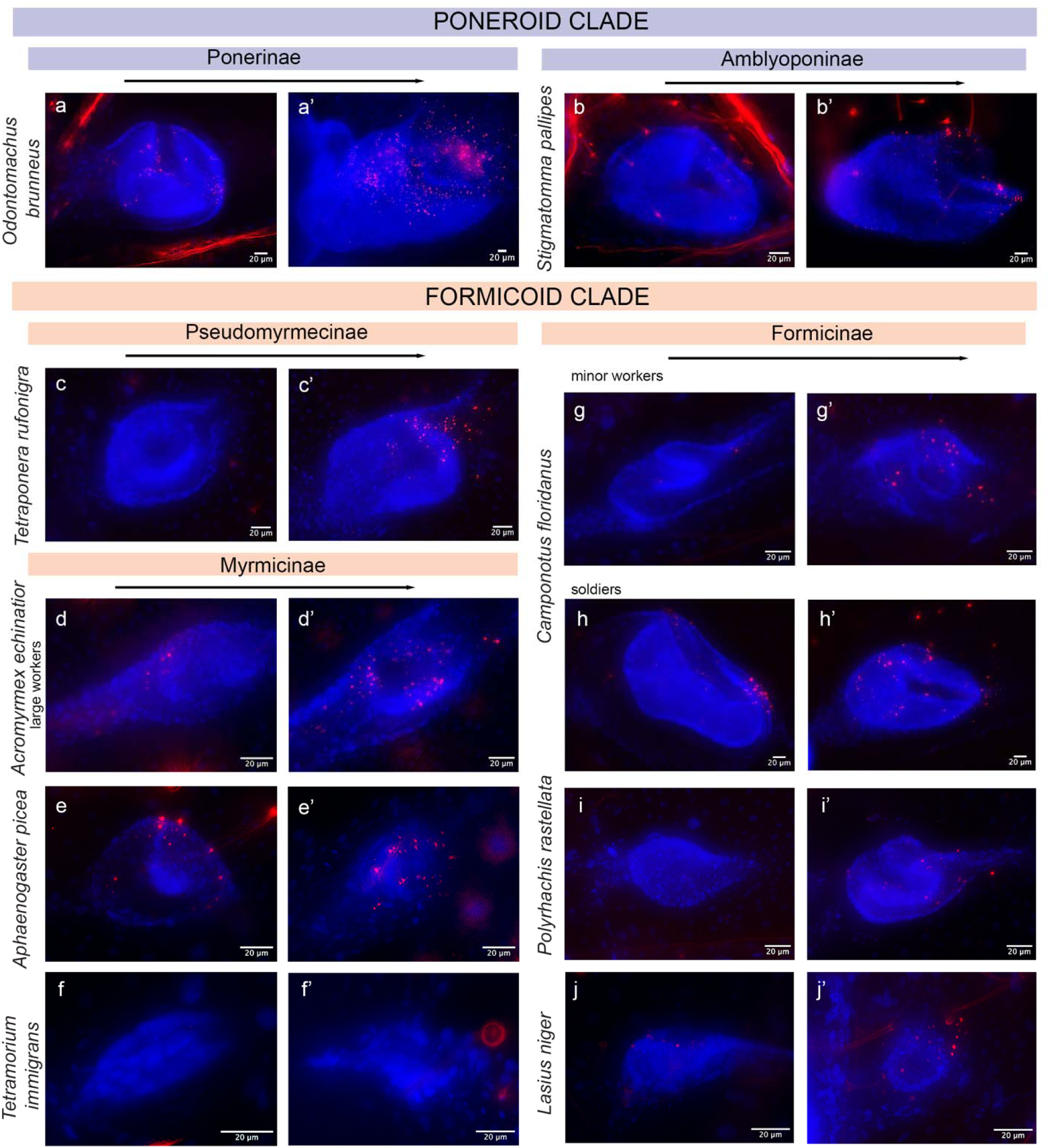
Programmed cell death in two stages of rudimentary wing disc development in larvae of wingless workers across the ant phylogeny. DAPI (blue) and TUNEL (red) signal in rudimentary wing discs of worker larvae from two species in the poneroid clade (a–b’) and eight species in the formicoid clade (c–j’). For each species, a rudimentary wing disc from the early to mid-terminal stage (left) and a rudimentary wing disc from the larval-prepupal transition stage (right) are shown. Black arrows indicate developmental time. Subfamily labels are marked on the top of the panels and species names are marked on the left of the panels. All wing discs are oriented with distal axis on the left and proximal axis on the right (see Fig. 1b).

We use the TUNEL assay as a general marker for apoptotic cell death (Kyrylkova et al., 2012). However, it remains possible that TUNEL may also detect other forms of PCD, such as autophagy (Grasl-Kraupp et al., 1995). We therefore used the active form of caspase 3, which has been shown to execute apoptosis (Xu et al., 2009), to confirm whether PCD is indeed carried out via the apoptosis pathway. Furthermore, we also tested for the activation of the autophagy pathway by using *atg8* and LC3A/B, genes known to be critical for autophagsome formation (Nakatogawa et al., 2007; Tanida et al., 2008). We found strong cleaved-caspase 3 expression at similar stages and regions within the rudimentary wing discs to the TUNEL assay in workers of three species belonging to the poneroid and formicoid clades: *H. saltator* (Ponerinae) (Fig. 4a, and a’), *Pheidole dentata* (Myrmicinae) (Fig. 4c’), and *Camponotus floridanus* (Formicinae) (Fig. 4e, and e’). In contrast, we found no detectable signal of either *atg8* and LC3A/B in the rudimentary wing discs of *H. saltator* and *P. dentata* workers, respectively (Fig. 4b, b’, d, and d’). Together these results confirm that PCD in the rudimentary wing discs of workers from various species across the ant phylogeny is carried out via the apoptosis pathway.

**Figure 4.**
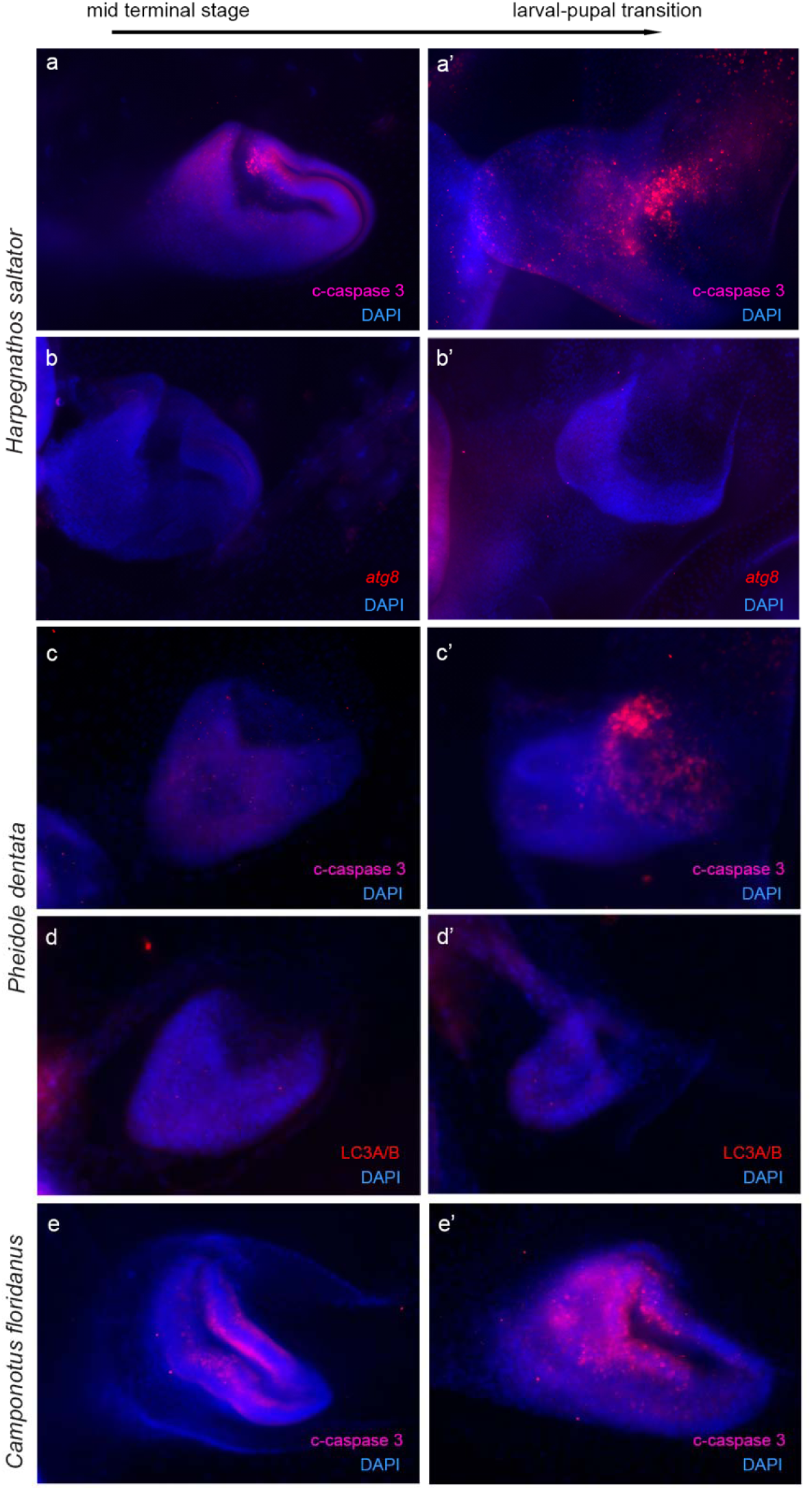
Programmed cell death is carried out via the apoptosis pathway. Expression of DAPI (blue) and cleaved-caspase 3 (pink) during mid (left) and late (right) terminal larval stages in *H. saltator* (a–a’), and *P. dentata* (c–c’), and soldier larvae of *C. floridanus* (e–e’). Expression of DAPI (blue) and autophagy marker *atg8* or LC3A/B (red) in rudimentary wing discs of *H. saltator* (b–b’) and soldier larvae of *P. dentata* (d–d’).

Finally, we performed an ancestral state estimation analysis to infer whether apoptosis was present or absent in the rudimentary wing discs of workers in the last common ancestor of ants. Our ancestral state estimation (under the equal rates model), using data from all species tested, strongly supports the presence of PCD (>95% confidence) as being ancestral to all living ants. This includes the poneroformicine clade (= poneroids + formicoids) (Fig. 5). Our inference remains robust after accounting for: (1) uncertainty in the presence or absence of PCD in species from missing ant subfamilies; (2) the absence of data for *Martialis* and Leptanillinae (the sister group to all other Formicidae); and (3) different taxon sampling regimes (Supp. 2). Altogether, our experimental data and ancestral state estimation analyses infer a role for PCD in the regulation of wing polyphenism in the last common ancestor of all extant ants (Fig. 5).

**Figure 5.**
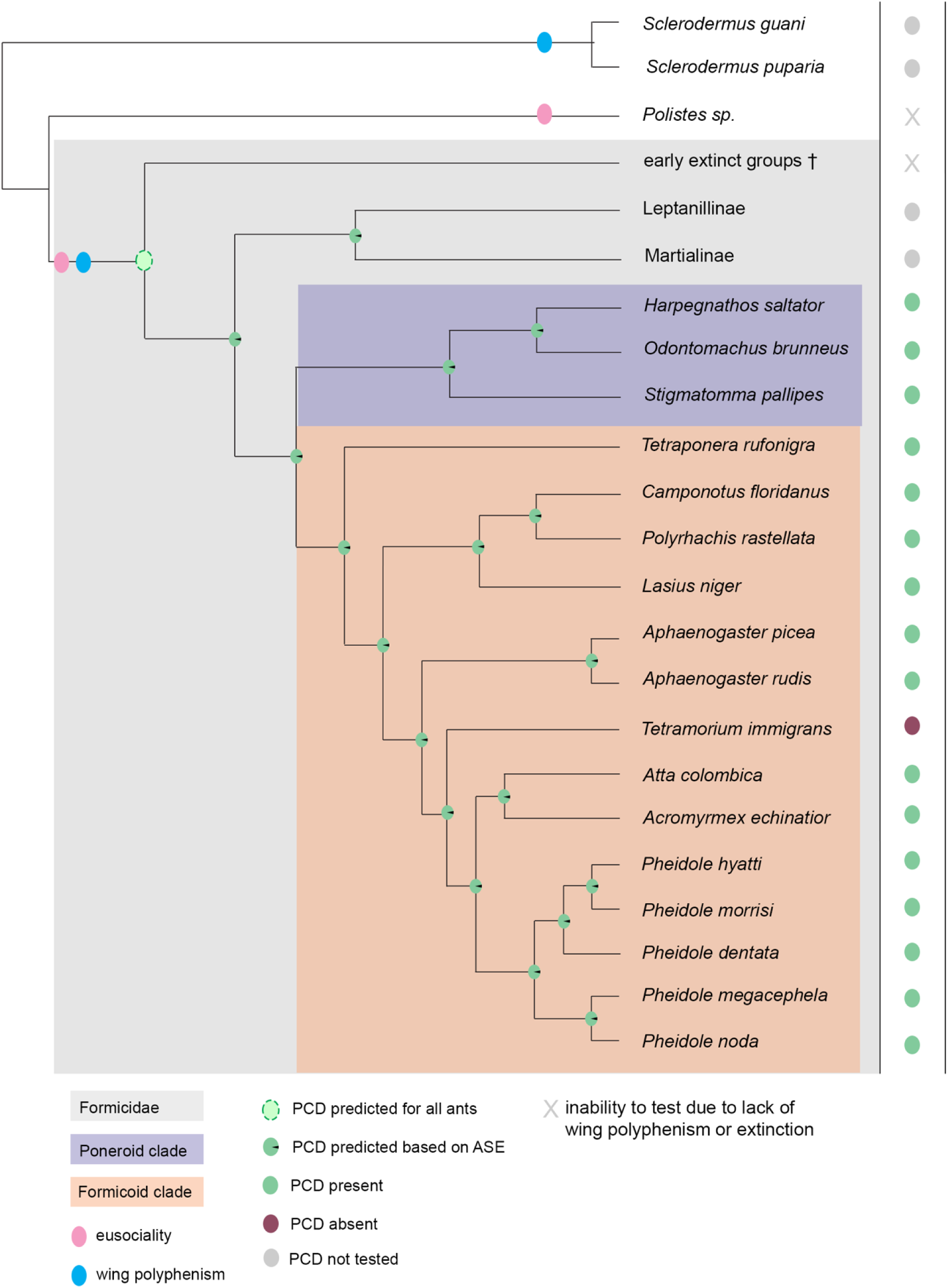
Programmed cell death in wing polyphenism across the ant phylogeny. Wing polyphenism (blue circle) and eusociality (pink circle) evolved concomitantly in ants (Formicidae, gray large box). Wing polyphenism and eusociality are found in some wasp species/groups. Outgroups for Formicidae are for demonstration purposes only and many branches in the Hymenoptera are excluded. Summary of PCD presence/absence found by this study for poneroid species (purple box) and formicoid species (orange box) are listed on the right. Characters indicate species with: presence of PCD (green circle), absence of PCD (red circle), PCD untested (gray circles). Gray X indicates species where testing is not possible. PCD regulation of wing polyphenism is predicted for all extant ants based on Ancestral State Estimations (ASE; green circles in nodes). ASE values for all tested models yielded over 95% confidence. PCD regulation of wing polyphenism at the origin of ants is proposed (dashed green circle). Branch length do not indicate evolutionary time. Phylogenetic relationships within the Formicidae are based on Boudinot, Khouri, et al. (2022); Moreau (2008); Ward et al. (2016).

## DISCUSSION

PCD has been proposed to play a role in major evolutionary transitions in the history of life, however, the examples used to support this hypothesis are limited to a few model species that do not necessarily reflect ancestral characteristics of the taxonomic groups they belong (Durand et al., 2019). We therefore used a phylogenetic comparative approach using 16 species spanning 5 of the major subfamilies of ants reflecting both ancestral and derived characteristics to formally test this proposal. Our analyses infer that PCD likely played a role in the origin of wing polyphenism, and therefore, in the major evolutionary transition to eusociality in ants. By helping to generate a wingless worker caste, PCD may have been critical in reinforcing the reproductive division of labor in ancestral ant societies by limiting dispersal of workers, and therefore, their participation in mating flights. Multiple ecological, evolutionary, and developmental factors influenced the evolution of eusociality in ants and in other eusocial organisms. Here, we show that PCD may have been one of these factors during the origin of eusociality in ants.

The occurrence of wing polyphenism in females has been recently confirmed to be present among the extinct stem lineages of ants (Boudinot, Khouri, et al., 2022), supporting PCDs role in wing polyphenism as a novel feature for all Formicidae, extant and extinct (Fig. 5 dotted green circle). Indeed, our analyses inferred that PCD is present and operating in the ‘basal-most’ subfamilies Leptanillinae and Martialinae; see, e.g., Romiguier et al. (2022), for which wing polyphenism in females is also known to occur (Fig. 5 green circles) (Borowiec et al., 2011; Chen et al., 2017; Hsu et al., 2017). However, empirically testing this inference in Leptanillinae is challenging given their sparse global distribution across Southeast Asia, Africa, and Western Europe. Testing this inference in Martialinae, however, is nearly impossible given that only two workers and a few males have been collected to date (Boudinot, 2015; Rabeling et al., 2008). Future developmental studies focusing on species within the Leptanillinae should further confirm PCD’s role in the origin of wing polyphenism in ants.

Many questions remain regarding how and from where PCD gained a regulatory role in the origin of wing polyphenism. Ants evolved from winged solitary wasps. However, in some of the wasp taxa closely related to the ants, a sexual wing dimorphism evolved multiple times independently, where the females are wingless or short winged and the males are fully winged (Hanna & Abouheif, 2021). Hanna and Abouheif (2021) proposed a “cryptic threshold model,” in which the winged solitary wasp ancestor from which the ants evolved may have retained a cryptic threshold to produce a wingless female morph. This model explains developmentally how a cryptic threshold could have been reactivated and repurposed in the ancestor of all ants to originate a female wing polyphenism. Furthermore, a recent review by Hanna and Abouheif (2023) showed that PCD has been co-opted repeatedly across the tree of life to facilitate the generation of alternative phenotypes, including sexual dimorphisms and polyphenisms. Based on this, we predict that PCD was likely operating to generate the sexual wing dimorphism in wasps, and reactivation of the cryptic threshold also reactivated PCD to facilitate the origin of wing polyphenism in ancestor of all ants.

Investigating the developmental mechanisms that coordinated with PCD to facilitate the origin of wing polyphenism in ants should be the focus of future studies. Key hormones, like JH and ecdysone, are sensitive to environmental and social cues such as nutrients, temperature, and pheromones (Edgar, 2006; Libbrecht et al., 2013; Wu & Brown, 2006). Therefore, changes in hormone levels or in the tissue sensitivity to these hormones may have facilitated the induction of PCD in the wing imaginal discs of select female individuals, reactivating the cryptic potential to generate a wingless female (giving rise to the wingless female caste in ants). Although currently there is no direct evidence for the role of hormones on regulating PCD pathways in ants, several studies in moths and beetles show a regulatory link between ecdysone hormone and the induction of apoptosis (Kijimoto et al., 2010; Lobbia et al., 2003; Niitsu et al., 2008). In addition, the insulin signaling pathway has been shown to be differentially expressed across ants (Chandra et al., 2018; Okada et al., 2010), and likely interacted with these hormones. Finally, investigating whether interruptions in the wing GRN in the worker castes of basal lineages are present and whether they interacted with PCD, hormones, and the insulin signaling pathway to regulate wing polyphenism will be key for understanding the origins or wing polyphenism and eusociality in ants.

To conclude, although it may not be immediately intuitive, we show that a destructive force like PCD at the level of the individual may have facilitated the origins of more complex division of labour at the colony-level. We present comparative evidence that PCD may have facilitated the origin of wing polyphenism and therefore eusociality in ants. By doing so, this raises the possibility that future studies using rigorous comparative approaches in other organisms across the tree of life may reveal that PCD facilitates major evolutionary transitions at different scales of biological organization, from multicellularity to eusociality.

## ACKNOWLEDGMENTS

We thank Llyod Davis, Robert Johnson, Luc Passera, and Angelly Vasquez-Correa for collecting ants. We also thank members of the Abouheif lab for comments on the manuscript. This work was supported by a Natural Sciences and Engineering Research Council (NSERC) Discovery Grant to E.A., and a Doctoral Fellowship from Fonds de Recherche du Quebec-Nature et Technologies (FRQNT) to L.H.

## MATERIALS AND METHODS

### Ant collection and colony care

Ant colonies were housed in plastic boxes lined with either fluon or talc powder. Artificial nests were constructed using glass test tubes half-filled with water and plugged with cotton. All colonies were maintained at 25 °C, 70% humidity and a 12h day:night cycle. Ants were fed a combination of mealworms, crickets, fruit flies, fruits, and Bhatkar-Whitcomb diet (Bhatkar & Whitcomb, 1970). *Harpegnathos saltator* colonies were housed in plaster nests and fed live crickets only 2-3 times per week.

Ant colonies were obtained from the following locals. *Stigmatomma pallipes* were collected at McGill Gault Nature Reserve (Quebec, Canada). *Lasius niger, Tetramorium immigrans, Aphaenogaster rudis*, and *Aphaenogaster picea* were collected at Mont Royal Park and McGill Gault Nature Reserve (Quebec, Canada). *Odontomachus brunneus* and *Camponotus floridanus* were collected at Gainesville (Florida, USA). *Pheidole dentata* were collected at Gainesville (Florida, USA) and Austin (Texas, USA). *Pheidole hyatti* were collected from Tempe (Arizona, USA). *Pheidole pallidula* were collected from Lyon (France). *Pheidole noda*, *Tetraponera rufonigra,* and *Polyrhachis rastellata* were collected from Mae Tang (Chiang Mai, Thailand). *Atta colombica* and *Acromyrmex echinatior* were collected from Gamboa (Panama). *Harpegnathos saltator* colonies were propagated in the laboratory since the original collection in 1999 from different locations in India.

### Immunohistochemistry, hybridization chain reaction, and TUNEL stainings

#### Fixation and Dissection

Terminal stage larvae were collected and fixed as previously described by Shbailat and Abouheif (2013). Fixed larvae were dissected using a Zeiss Discovery V12 stereomicroscope to expose the wing discs and remove any obstructing fat tissue.

#### Immunohistochemistry

The following primary antibodies were used for immunohistochemistry: anti-cleaved caspase-3 (1:100-1:200, Cell Signaling Technology, #9661) and anti-LC3A/B (1:100, Cell Signaling Technology, #4108). For all immunohistochemistry assays, fluorescent secondary anti-rabbit polyclonal Alexa Fluor-555 (AbCam) antibody was used at 1:500 dilution to detect the primary antibody, according to Khila and Abouheif (2008).

#### Hybridization Chain Reaction

To detect mRNA expression of *H. saltator atg8* we utilized the hybridization chain reaction (HCR) methodology (Choi et al., 2018). Probe sets, amplifiers, and buffers were purchased from Molecular Instruments, Inc. Procedure followed based on the Protocols for HCR^TM^ RNA-FISH (v3.0) (generic sample in solution) acquired from molecularinstruments.com. Prior to procedure, samples were fixed and dissected as described above, rehydrated in 25%, 50%, 75% and 100% PTw (1X PBS; 0.1% Tween 20) and permeabilized in PTw and 2% Triton X-100 for 1 hour. Larvae were post-fixed in 4% formaldehyde in SSCT (5X sodium chloride sodium citrate; 0.1% Tween 20) for 2 hours and then washed 3 times with SSCT before moving into glycerol.

### TUNEL

To detect programmed cell death in the wing discs of all species, we used the TUNEL (Terminal deoxynucleotidyl transferase dUTP nick end labeling) assay. We used the *In Situ* Cell Death Detection Kit, TMR red (Roche). Larval samples were fixed, dissected and stored in 100% methanol in −30 °C prior to procedure. Dissected larvae were then rehydrated in 25%, 50%, 75%, and 100% PTw (1X PBS; 0.1% Tween 20). Samples were then washed 3 times (10 min each) in freshly made PBT (1X PBS; 0.1% Triton; 0.1% BSA) and permeabilized in PTw and 2% Triton X-100 for 1 hour, followed by 5 min washes in PBT. Positive and negative controls were incubated in DNase buffer at 37°C for 10 minutes and subsequently washed with PBT for 5 min (Supp. 3). Enzyme mix and label solution were added to the samples and the positive controls (only label solution was added to negative controls) and incubated in the dark at 37 °C for 1 hour followed by 3 washes in PTw 5 min each. Samples were stained with DAPI (1:1000) for 1 hour to overnight and followed by gradual moving into glycerol. Samples were stored in final concentration of 85% glycerol/DAPI before mounting and imaging.

#### Imaging

Fluorescent images of larval rudimentary wing discs of all species utilized in this study were taken using a Zeiss AxioImager Z1 microscope.

#### Ancestral state estimation

To reconstruct the ancestral conditions of PCD, we conducted a series of ancestral state estimation (ASEs) analyses given prior and new experimental data. To ensure that our estimates were based on well-supported phylogenetic results, we used the favored phylogram (P) and chronogram (C) of Romiguier et al. (2022), particularly as a recent simulation study has demonstrated the importance of branch lengths for ancestral state modeling (Wilson et al., 2022). We pruned the two trees (P, C) to three alternative taxon sets: (P1, C1) the first included only those terminals that match species with experimental data (17 taxa), (P2, C2) the second included experimental terminals plus *Martialis* and a leptanilline (22 taxa), and (P3, C3) the third included all ant terminals from the Romiguier et al. (2022) dataset in order to account for phylogenetic uncertainty (74 taxa). Because we had surplus experimental sampling of *Aphaenogaster* and *Pheidole* with identical results, we included only one species of the former and two species of the latter, as this matched the phylogenomic sampling of Romiguier et al. (2022). We also excluded non-ant outgroups as there is no applicable experimental data, thus insufficient information to parameterize a phylogenetic model that includes these taxa. As the ‘all rates different’ model overfit the data, resulting in complete statistical separation, we only report results under the ‘equal rates’ model.

---

**Supplemental 1.**
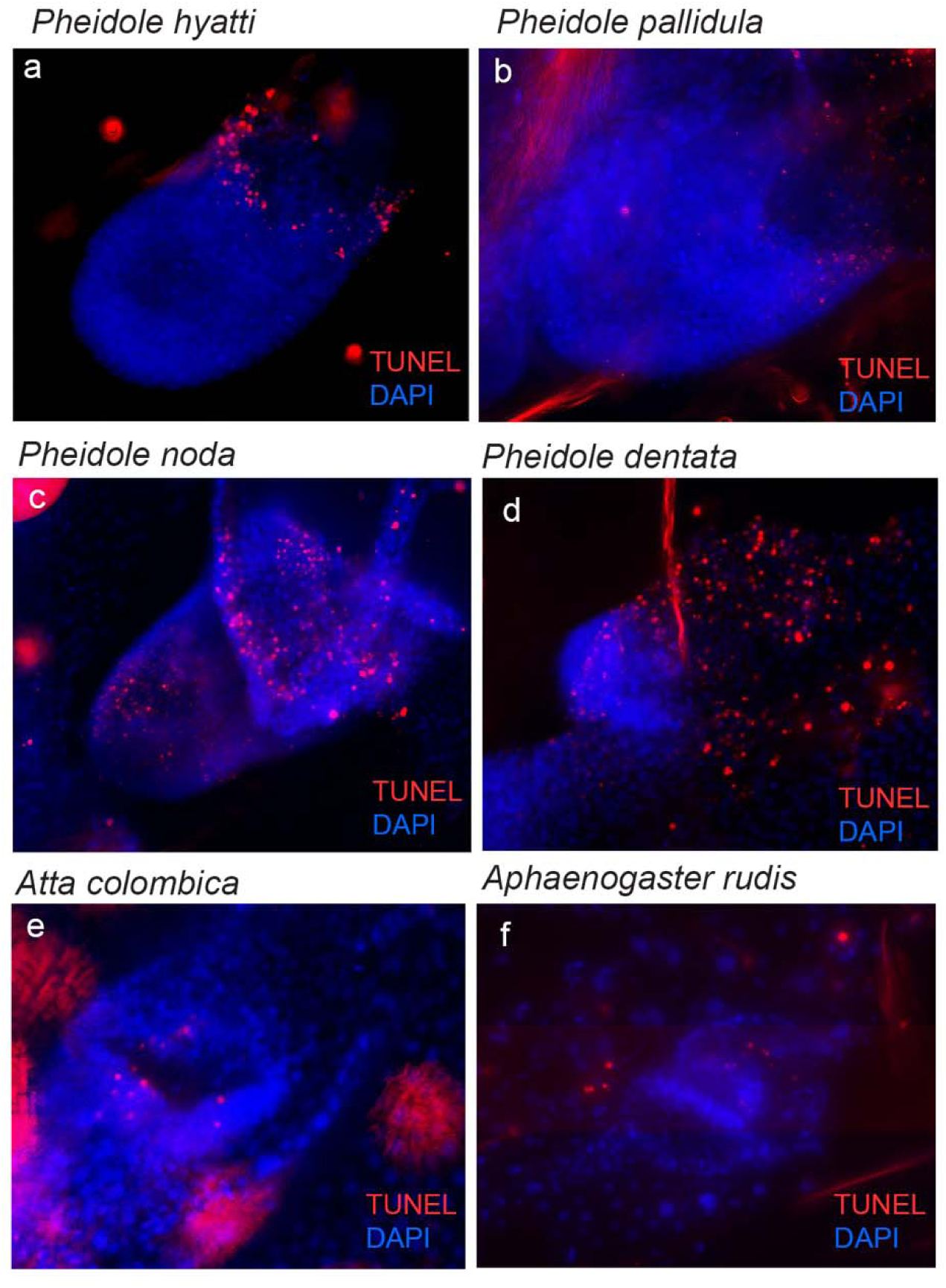
Programmed cell death in additional ant species from the subfamily Myrmicinae. For all panels, a rudimentary wing discs at the late terminal larval stage is shown stained in DAPI (blue) and TUNEL (red).

**Supplemental 2.**
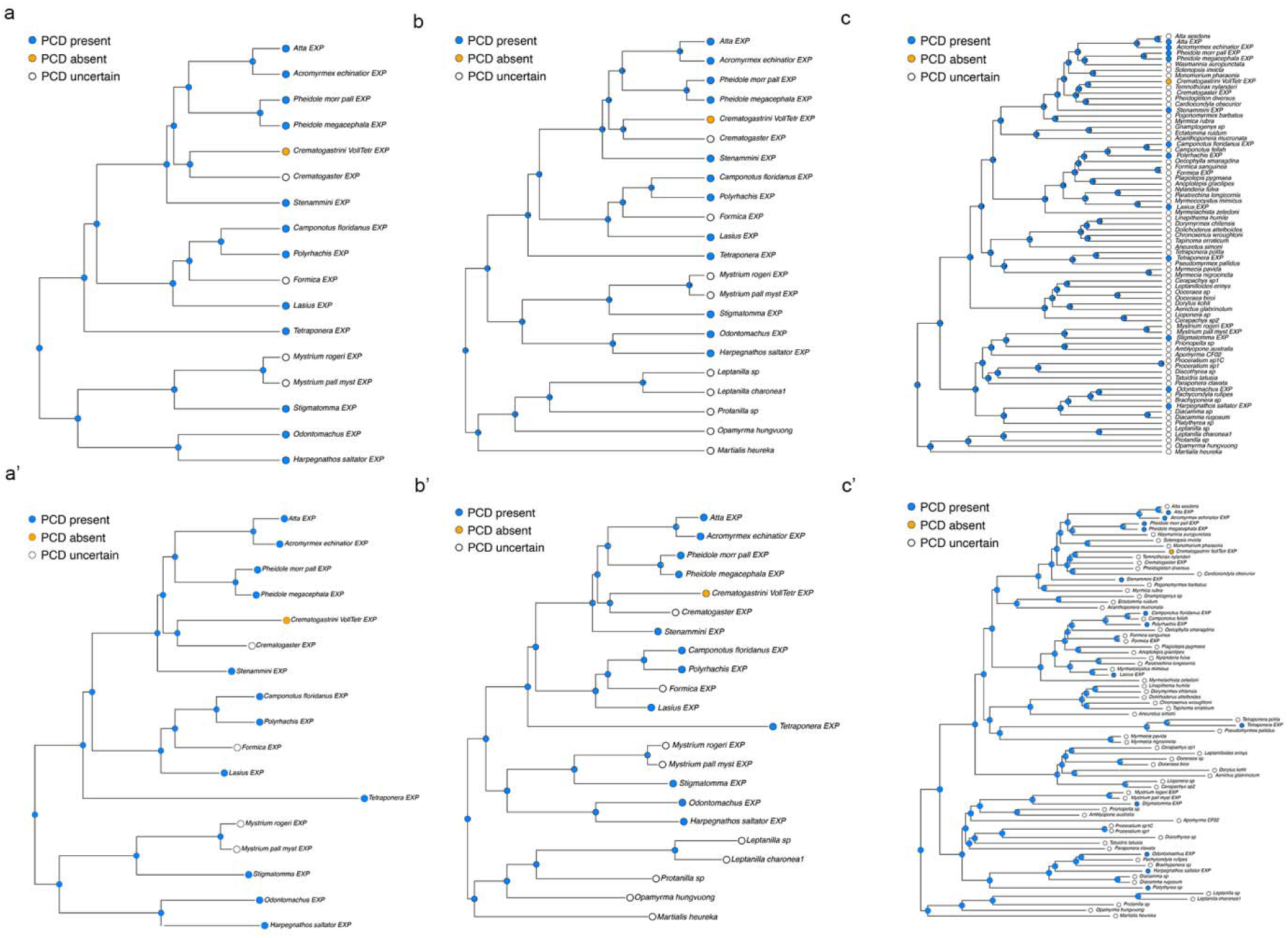
Ancestral State Estimations for presence and absence of PCD. Estimates based only on species where assays for PCD have been performed are shown using a phylogram (a) and chronogram (a’) modeling. Estimates based on species where assays for PCD have been performed as well as the addition of basal outgroups *Martialis* and leptanilline are shown using a phylogram (b) and chronogram (b’) modeling. Estimates based on 74 ant taxa including those without experimental data are shown using a phylogram (c) and chronogram (c’) modeling. Filled blue circles indicate estimates for presence of PCD; open circles indicate estimates for absence of PCD, open circle indicate species where PCD has not been examined. All models yielded over 95% confidence.

**Supplemental 3.**
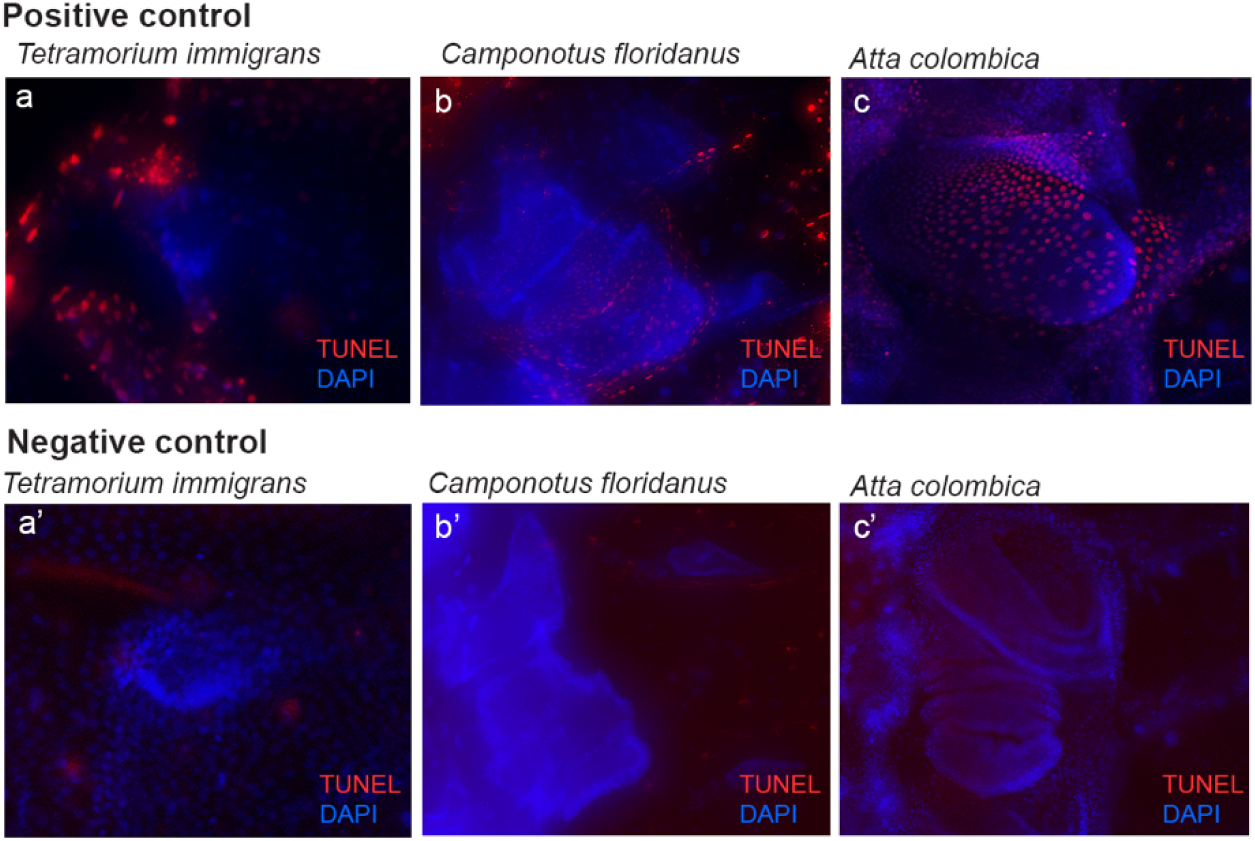
Positive and negative TUNEL controls shown for three ant species. DAPI (blue), TUNEL (red).

